# Semisynthesis of Oxalyl-Coenzyme A for Enzymatic Assays

**DOI:** 10.64898/2026.08.06.743301

**Authors:** Sergey Nepogodiev, Martin Rejzek, Megan N. Steinberg, Anne Edwards, Cathie Martin

## Abstract

Oxalyl-coenzyme A (oxalyl-CoA) is a key intermediate in oxalate metabolism in plants, fungi and oxalate-degrading bacteria, but its limited availability has restricted biochemical investigations of oxalyl-CoA-dependent enzymes. Here, we describe a practical semisynthetic procedure for the preparation of oxalyl-CoA based on rapid oxalyl transfer from *S*-oxalyl *p*-thiocresol to coenzyme A. The reaction was monitored directly by ^1^H NMR spectroscopy, allowing optimisation of pD and reaction conditions. Following removal of thiocresol and purification by reversed-phase HPLC, oxalyl-CoA was obtained in 39% yield as determined by quantitative ^1^H NMR. The product was characterised by high-resolution electrospray mass spectrometry and comprehensive ^1^H, ^13^C and ^31^P NMR spectroscopy, confirming its structure unequivocally. During the study, the limited stability of oxalyl-CoA in aqueous solution was documented, leading to recommendations for its purification and storage. The semisynthetic protocol provides a convenient source of analytically pure oxalyl-CoA suitable for biochemical assays and supplies reference spectroscopic data for its unambiguous identification. The biological utility of the semisynthetic oxalyl-CoA was demonstrated by its application as an acyl donor substrate in assays of PnBAHD15, enabling quantitative kinetic characterisation of the enzyme and illustrating its suitability for biochemical studies of oxalyl-CoA-dependent enzymes.

## Introduction

Oxalyl-coenzyme A (oxalyl-CoA) is a central intermediate in oxalate degradation in plants, fungi and many microorganisms. In yeast, degradation of oxalate via oxalyl-CoA is thought to be the only pathway in which oxalyl CoA serves as an intermediate. Oxalyl-CoA is an important intermediate in plant defence against oxalate-producing fungal pathogens like *Sclerotinia* spp. Several intestinal bacteria, including *Oxalobacter formigenes* and species of *Bifidobacterium*, use oxalyl-CoA pathways to extract energy from dietary oxalates. Humans ingest significant amounts of oxalates through foods like spinach, tea, chocolate, and coffee but oxalate itself is toxic and excess oxalate can cause the formation of calcium oxalate, hyperoxaluria and kidney stones. Humans lack an endogenous pathway for oxalate degradation and therefore rely largely on the intestinal microbiota to break oxalate down. Within this pathway, oxalyl-CoA is decarboxylated by oxalyl-CoA decarboxylase to formyl-CoA and carbon dioxide. Oxalyl-CoA decarboxylase is highly specific for its substrate, oxalyl-CoA, making it an important intermediate for many analyses of detoxification pathways.^1^ Oxalyl-CoA is highly reactive, allowing it to act as an acyl donor in numerous biochemical reactions but also rendering it difficult to prepare by enzymic synthesis. Consequently, most assays of the acyl-activating enzyme AAE3 that synthesises oxalyl-CoA from oxalate and CoA in plants and fungi, rely on a coupled assay that measures the breakdown of ATP to AMP as a proxy for oxalyl-CoA because of difficulties associated with purification of oxalyl-CoA.^2,3,4^

To address the need for authentic oxalyl-CoA standards and substrates for biochemical assays, we synthesised oxalyl-CoA chemically using the method originally described by Quayle in 1962.^5^ The method utilises high reactivity of oxalyl thioesters toward thiols and takes advantage of relative stability and water solubility of *S*-oxalyl thiocresol used as the acylating reagent. Despite various reports of preparation of CoA thioesters,^6^ Quayle’s method remains superior for oxalyl-CoA. Thus, a more recent Chinese patent^7^ describes a variant of the method, but provides little detail regarding purification and analytical characterisation of the oxalyl-CoA produced. Here we report a practical protocol for the semisynthesis, HPLC purification, HRMS and NMR characterisation of oxalyl-CoA. The resulting material is suitable for direct biochemical assays of oxalyl-CoA-dependent enzymes as demonstrated by enzymatic assays of purified PnBAHD15, an enzyme from *Panax notoginseng* that can synthesise dencichine also known as β-*N*-oxalyl-L-α,β-diaminopropionic acid (β-ODAP). Using oxalyl-CoA and L-2,3-diaminopropionic acid (L-DAP) as substrates, we evaluated β-ODAP synthesis and determined the enzyme’s kinetic parameters toward oxalyl-CoA.

## Results and discussion

### Synthesis, purification and characterisation of oxalyl-CoA

Chemical preparation of CoA thioesters is best regarded as semisynthesis because it relies on derivatisation of the naturally occurring coenzyme A.^8^ Various *S*-acylating reagents have been employed for the preparation of CoA thioesters.^6^ For oxalyl-CoA, *S*-oxalyl thiocresol proved to be particularly suitable because it readily transfers the oxalyl group to thiols.^9^ This reagent can be readily prepared by general method of Stolle^10^ by the reaction of *p*-thiocresol (**1**) with excess oxalyl chloride. Following hydrolysis of residual acyl chlorides, aqueous work-up and silica gel chromatography, thiooxolate **2** was obtained as a white solid in 75% yield (Scheme 1).

The reaction of CoA (**1**) with thiooxolate **2** was deliberately performed in an NMR tube to permit real-time monitoring of product formation and optimisation of pD without sample transfer. To proceed, the thioesterification required weakly basic conditions. Consequently, NaOD was added incrementally to maintain the reaction while pD was checked periodically. The reaction reached completion at approximately pD 7.5 as judged by the absence of further changes in the ^1^H NMR spectra (Fig. 1A & 1B). The majority of *p*-tiocresol that was poorly soluble in water was released from reagent **2** and removed by extraction with EtOAc (Fig. 1C). Finally, reverse-phase HPLC under conditions suitable for purification of CoA derivatives (Supplementary Fig. 1) provided pure oxalyl-CoA (Fig. 1D). The isolated yield (39%) of oxalyl-CoA was determined by quantitative NMR rather than gravimetrically (Supplementary Fig. 2).

**Figure 1.**
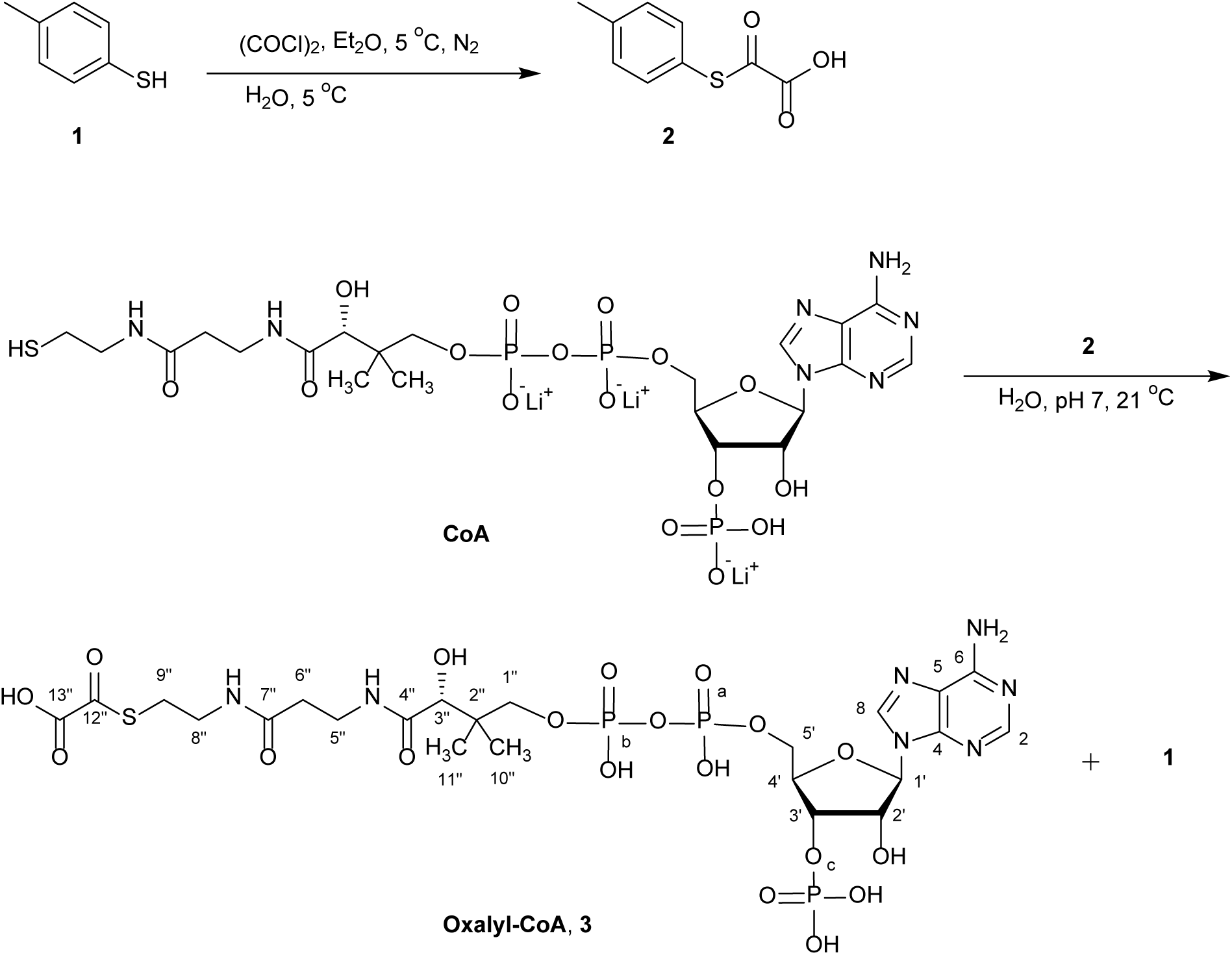
Synthesis of oxalyl-CoA.

**Figure 1.**
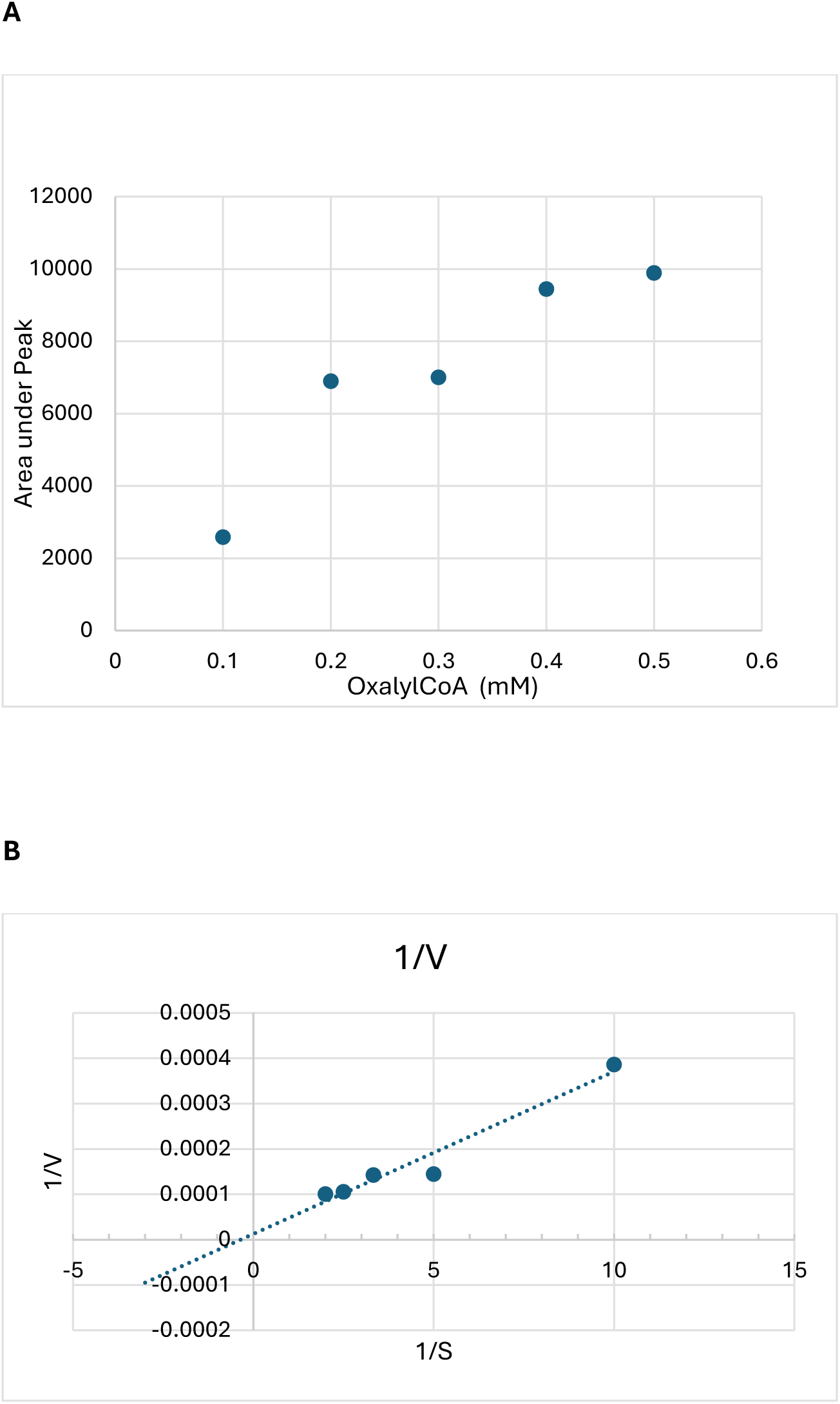
Enzyme assays using synthesised oxalyl-CoA as a substrate to make β-ODAP. *Pn*BAHD15 from *Panax notoginseng* was expressed in *E. coli* and purified. (**A**) oxalyl CoA production in response to increasing concentrations of synthetic oxalyl CoA with a fixed amount of enzyme. (**B**) Plot of 1/velocity against 1/[oxalyl CoA in mM] to determine the Km of *Pn*BAHD15 for oxalyl CoA. Assays allowed the Km for oxalyl CoA to be calculated as 250 μM.

The structure of synthesised compound **3** was confirmed by HR–ESI–MS and NMR spectroscopy analyses. Mass spectra of **3** were measured in negative mode and gave the expected pseudomolecular ion of oxalyl-CoA along with a number of Na^+^ adducts (Table 1, Supplemetary Fig. 3). The most intense ions in the ESI–MS of **3** corresponded to cleavage of the oxalyl thioester bond, producing CoA ions assigned as [M−2H]^2−^ and [M−H]^−^.

**Table 1.** Assignment of HR–ESI–MS ions observed for oxalyl-CoA.

| Peak | *m/z*<br>(observed) | Ion<br>assignment | *m/z*<br>(calculated) | Error<br>(ppm) | Intensity |
| --- | --- | --- | --- | --- | --- |
| 1 | 382.5508 | CoA [M-2H]<sup>2-</sup> | 382.5508 | 1.3 | 100 |
| 2 | 766.1083 | CoA [M-H]<sup>-</sup> | 766.1079 | 0.5 | 24 |
| 3 | 788.0895 | CoA [M+Na-2H]<sup>-</sup> | 788.0899 | 0.5 | 13 |
| 4 | 810.0717 | CoA [M+2Na-3H]<sup>-</sup> | 810.0718 | 0.1 | 7.5 |
| 5 | 838.0932 | Oxalyl-CoA [M-H]<sup>-</sup> | 838.0927 | 0.6 | 13 |
| 6 | 860.0744 | Oxalyl-CoA [M+Na-2H]<sup>-</sup> | 860.0746 | 0.2 | 6 |
| 7 | 882.0562 | Oxalyl-CoA [M+2Na-2H]<sup>-</sup> | 882.0566 | 0.4 | 4 |

Assignment of ^1^H and ^13^C and ^31^P NMR spectra of **3** was achieved using 2D ^1^H,^1^H-COSY, ^1^H,^13^C-HSQC, ^1^H,^13^C-HMBC and ^1^H,^31^P-HMBC experiments (see Supplementary Figs. 4−11), and the resulting assignments are summarised in Table 2. Presence of the oxalyl group in compound **3** was confirmed by two characteristic resonances in the ^13^C NMR spectrum at δ 194.8 (thiocarbonyl carbon, C12”) and δ 164.4 (carboxyl carbon, “C13”). Acylation of the CoA thiol was confirmed by the strong HMBC correlation between methylene protons H9” (CH_2_S) and C12”, and by the characteristic downfield shift of H9” resulting from deshielding by the oxalyl group. Assignment of phosphorous resonances was achieved from the well-resolved^1^H,^31^P-HMBC cross-peaks H3’/P-c, H5’/P-a and H1”/P-b (Supplementary Fig. 11).

**Table 2.** Resonance assignment in 13C, 1H and 31P NMR spectra of oxalyl-CoA.

| No | $\delta_{\text{C}}^*$ | $\delta_{\text{H}}$ (multiplicity, $J$) | $\delta_{\text{P}}$ |
| --- | --- | --- | --- |
| 2 | 151.1 | 8.29 (s) | — |
| 4 | 149.2 | — | — |
| 5 | 118.6 | — | — |
| 6 | 154.4 | — | — |
| 8 | 140.4 | 8.56 (s) | — |
| 1' | 86.5 | 6.18 (d, 6.6 Hz) | — |
| 2' | 74.2 (d, 5.1 Hz) | 4.87 (m) | — |
| 3' | 73.7 (d, 5.0 Hz) | 4.84 (m) | — |
| 4' | 83.5 (dd, 3.2 Hz; 9.2) | 4.59 (q, 2.7 Hz) | — |
| 5' | 65.2 (d, 5.3 Hz) | 4.25 (m) | — |
| 1'' | 71.8 (d, 6.0 Hz) | — | — |
| 1''b | — | 3.55 (dd, 4.6 Hz, 9.8 Hz) | — |
| 1''a | — | 3.88 (dd, 4.8 Hz, 9.8 Hz) | — |
| 2'' | 38.3 (d, 8.2 Hz) | — | — |
| 3'' | 74.1 | 4.01 (m) | — |
| 4'' | 174.7 | — | — |
| 5'' | 35.4 | 3.45 (m) | — |
| 6'' | 35.3 | 2.43 (t, 6.7 Hz) | — |
| 7'' | 174.0 | — | — |
| 8'' | 38.4 | 3.35 (t, 6.5 Hz) | — |
| 9'' | 28.1 | 3.02 (t, 6.5 Hz) | — |
| 10'' | 20.8 | 0.89 (s) | — |
| 11'' | 18.0 | 0.75 (s) | — |
| 12'' | 194.8 | — | — |
| 13'' | 164.4 | — | — |
| a | — | — | -11.4 (d, 21.0 Hz) |
| b | — | — | -10.8 (d, 21.0 Hz) |
| c | — | — | -0.25 (s) |

\* Multiplicities and $J$ coupling are given for resonances of splitted due to coupling with $^{31}\text{P}$ (C-2' – C5', C1'' and C-2'').

During NMR analysis we observed gradual decomposition of oxalyl-CoA, consistent with the known lability of CoA thioesters. Although CoA thioesters are sufficiently stable under mildly acidic to neutral conditions (pH 4–7), they remain susceptible to rapid hydrolysis, decarboxylation, or trans-thioesterification. Previous studies^11^ have shown that non-degassed NMR samples of CoA are prone to oxidation, forming disulfides. Indeed, from the changes of chemical shifts of H-9’’ and H-8’’ in the ^1^H NMR spectrum of a sample of CoA stored for three weeks in D_2_O in the fridge (4 °C), it was evident that CoA was completely converted into the disulfide. Chemical shifts of H9’’ and H8’’ similar to those of CoA disulfides were also observed in the ^1^H NMR spectrum of oxalyl-CoA after keeping its D_2_O solution in the fridge for seven days (Supplementary Fig. 12). That indicated hydrolysis of the thioester of oxalyl-CoA followed by even faster oxidation of the resulting CoA to the disulfide. Accordingly, purified fractions were freeze-dried immediately after HPLC purification and the resulting solid was stored at −18 °C.

### Testing the efficacy of semisynthesised oxalyl CoA

*Pn*BAHD15 is an enzyme that can react with *Pn*AAE3 (oxalyl CoA synthase) and use oxalic acid and *L*-DAP as substrates and CoA as a cofactor to produce β-ODAP.^12^ We performed *in vitro* experiments to investigate whether *Pn*BAHD15 could use semisynthesised oxalyl-CoA and *L*-DAP to produce β-ODAP directly. Our results showed that *Pn*BAHD15 could use oxalyl-CoA and *L*-DAP as substrates to produce β-ODAP directly. Kinetic analysis showed that the *K_m_* values of this reaction were 250 μM for oxalyl-CoA (Figs. A, B).

### Conclusions

A practical semisynthetic protocol for the preparation of oxalyl-coenzyme A was established using rapid oxalyl transfer from *S*-oxalyl *p*-thiocresol to coenzyme A. The reaction was conveniently monitored by ^1^H NMR spectroscopy, enabling optimisation of the reaction pD and purification procedure. The identity and purity of the product were confirmed by HR–ESI–MS together with complete ^1^H, ^13^C and ^31^P NMR assignments. Although oxalyl-CoA was found to be susceptible to decomposition in aqueous solution, immediate freeze-drying and storage at −18 °C provided material suitable for subsequent biochemical studies. The applicability of the semisynthetic oxalyl-CoA was demonstrated in enzymatic assays, illustrating its value as a substrate for investigations of oxalyl-CoA-dependent enzymes.

## Materials and Methods

*p*-Thiocresol, oxalyl chloride and Coenzyme A trilithium salt (MW 767.53) were purchased from Merck. For reactions carried out in NMR tubes, a manual centrifuge was used to sediment precipitates and separate the aqueous and organic phases. Sample purification was carried out using a Biotage Isolera medium-pressure purification system using 10 g silica gel cartridges and semi-preparative Thermo Dionex Ultimate 3000 HPLC equipped with UV detector (265 nm) set for detection at 265 nm on Phenomenex Kinetex XB–C18 reversed-phase column (250 × 21.2 mm, 5 μm, 100 Å). UPLC–ESI–HRMS analysis was performed using a Waters Acquity UPLC system equipped with a Micromass Q–TOF Premier mass spectrometer (Waters MS Technologies, Manchester, UK). NMR spectra were acquired on a Bruker Avance Neo 600 MHz spectrometer equipped with a 5 mm TCI cryoprobe, and on a Bruker Avance III 400 MHz spectrometer equipped with a broadband BBO probe. Data were processed with TopSpin 4.1 or MestReNova (Mnova) 15 software. Unless otherwise stated, NMR experiments were carried out in D2O at 298 K. Residual H_2_O signals in the ^1^H NMR spectra were suppressed using presaturation. Resonance assignments were based on analysis of 1D DEPT-135 and 2D ^1^H,^1^H-COSY, ^1^H,^13^C-HSQC-edited, ^1^H,^13^C-HMBCed and ^1^H,^31^P-HMBC spectra. pD of NMR samples was measured using Thermo Scientific Orion micro pH electrode.

### Caution

Thiocresol and the by-products generated during the synthesis of *S*-oxalyl thiocresol and oxalyl-CoA possess a strong, unpleasant odour. Waste should be treated with sodium hypochlorite solution (bleach) before disposal.

### *S*-Oxalyl *p*-thiocresol (2-Oxo-2-(4-methylphenylthio)acetic acid)

To a stirred solution of oxalyl chloride (0.7 mL, 8.2 mmol) in anhydrous Et_2_O (15 mL) cooled in an ice bath, was added dropwise a solution of thiocresol (700 mg, 5.7 mmol) in anhydrous Et_2_O (20 mL) under the slow stream of nitrogen. After 1 h, the ice bath was removed and the reaction mixture was stirred at room temperature for a further 19 h. Volatiles were removed by evaporation in vacuo, the residue was dissolved in EtOAc (20 mL), cooled in an ice bath, water (50 mL) was added cautiously and the biphasic mixture was stirred vigorously at room temperature for 1 h. The aqueous layer was separated and extracted with ethyl acetate (3 × 10 mL). The original organic phase and extracts were combined, dried over MgSO_4_, and concentrated to dryness on a rotary evaporator giving yellowish crude solid (970 mg). The crude product was purified by flash chromatography using 10 g silica gel cartridge eluting with a gradient of hexane–ethyl acetate (20–100% ethyl acetate). Removal of solvent under reduced pressure afforded the product as white crystals (840 mg, 75%). ^1^H NMR (400 MHz, CDCl_3_): δ 8.69 (s, 1H), 7.35–7.26 (m, 4H), 2.41 (s, 3H); ^13^C NMR (150 MHz, CDCl_3_): δ 186.9, 158.5, 141.0, 134.0, 130.6, 121.5, 21.4. The NMR data were in good agreement with published values.^13^

### Oxalyl-Coenzyme A (Oxalyl-CoA) (3)

Oxalyl-CoA was synthesised at 22 °C in D_2_O using a 5 mm NMR tube as the reaction vessel, allowing the reaction to be monitored directly by ^1^H NMR spectroscopy on a 400 MHz spectrometer. The reaction pD was pH-monitored using a microelectrode, and estimated as pH + 0.4. To a solution of coenzyme A (trilithium salt, 10.32 mg, 13 μmoles) in 300 μL D_2_O (which had an initial pD of 5.3) was added a solution of **1** (3.17 mg, 16 μmoles) in D_2_O (300 μL) resulting in a decrease in pD from 5.3 to 2.5. The solution pD was adjusted to 7.2–7.6 by incremental addition of 100 mM NaOD in D2O (28 μL total), with periodic vortex mixing. Prior to each intermediate ^1^H NMR measurements, the reaction mixture was centrifuged to sediment the precipitate and obtain a clear supernatant in the NMR tube. ^1^H NMR analysis confirmed conversion of CoA into oxalyl-CoA (**3**) (Fig. 2B). The suspension of the reaction mixture was extracted with EtOAc (2 × 0.25 mL) to remove poorly soluble thiocresol, yielding a clear aqueous phase. Removal of thiocresol was confirmed by repeated ^1^H NMR of D_2_O solution (Fig. 2C). The solution was loaded onto a C18 Sep-Pak cartridge and the product was eluted with H_2_O (2 mL). That solution was freeze-dried and the crude product was purified by reverse phase chromatography on Phenomenex Kinetex column (5 μm XB-C18, 100 Å, 250 x 21.2 mm) using solvent A (10 mM aqueous ammonium formate) and solvent B (acetonitrile). The column was eluted at 20 mL min^−1^ using the following gradient: 0–6 min, 100% A; 6–12 min, 0 to 10% B; 12–15 min 10% B (Supplementary Fig. 1A). Fractions collected between 11.2 and 12.1 min, corresponding to the major chromatographic peak (Supplementary Fig. 1B), were pooled and immediately freeze-dried. Freeze-dried samples of oxalyl-CoA were stored at −18 °C. Product purity was confirmed by ^1^H NMR (Fig. 2D). Purified oxalyl-CoA (3) was obtained in 39% yield (4.0 mg), as determined by quantitative ^1^H NMR using the TopSpin ERETIC protocol (Supplementary Fig. 2).

**Figure 2.**
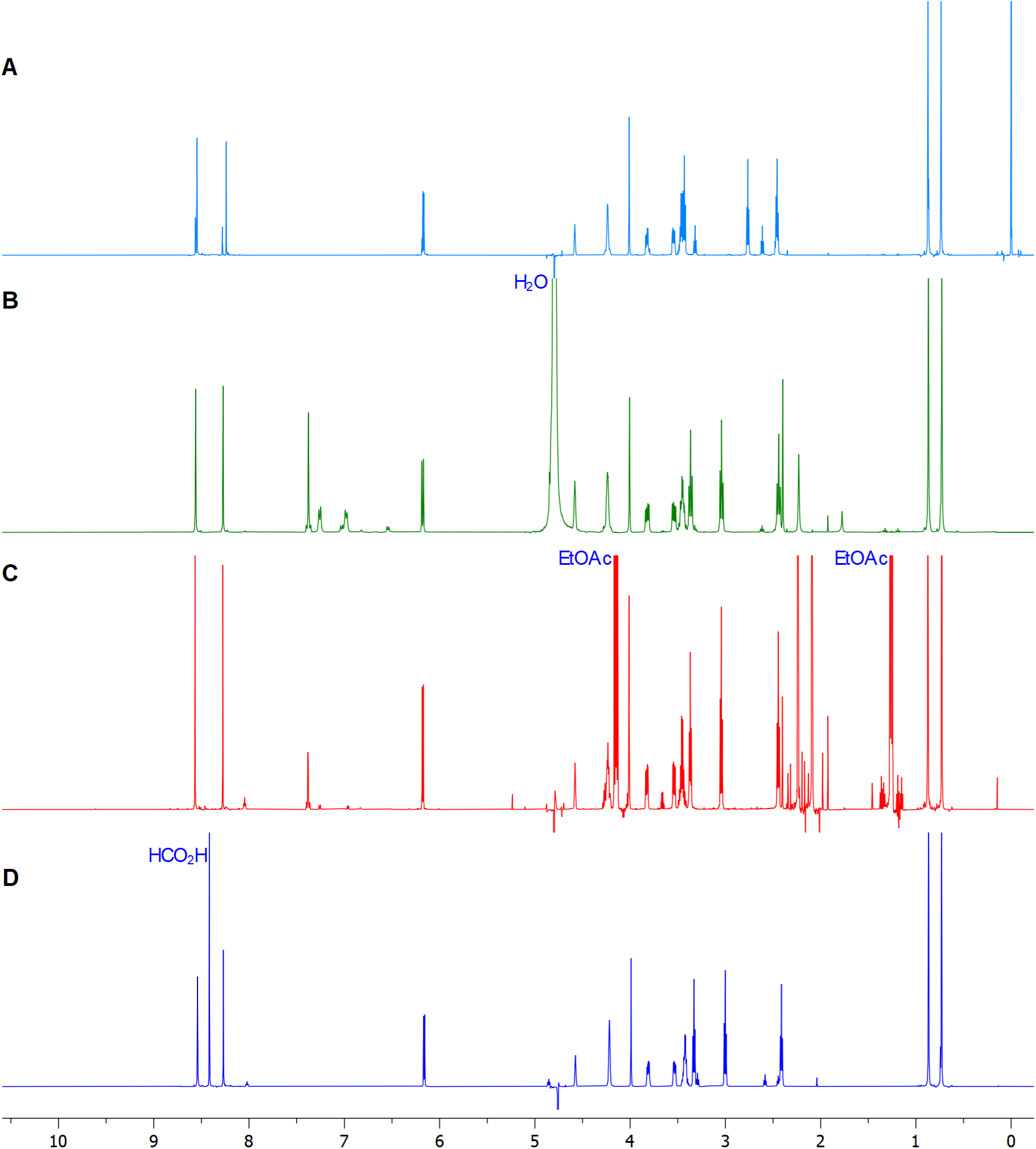
Stack of ^1^H NMR spectra showing progress of the formation of oxalyl-CoA. (**A**) pure commercial CoA in D_2_O phosphate buffer, pD 7.0; (**B**) Crude product **4** of the reaction carried out in D_2_O, pD 7.4; (**C**) Reaction mixture after extraction of thiocresol byproducts ed with EtOAc to remove; (**D**) Purified oxalyl-CoA (4) in D_2_O, pD 7.2. All ^1^H spectra except (**B**) were recorded with water suppression (Bruker noesypresat pulse program). Signal at 8.4 ppm in (**D**) is due to the formate captured from theHPLC solvent.

### Characterisation of oxalyl-CoA by mass-spectrometry and NMR spectroscopy

ESI–HRMS and NMR analysis were conducted using instrumentation described in the General Methods and the resulting spectra are shown in Supplementary Figs. 3–11.

### Expression of PnBAHD15 and protein purification

For *E. coli* expression, the amplified PnBAHD15 cDNA was cloned into pDEST17 (N terminal His tag expression vector; Lifetech), transformed into Arctic Express (DE3) competent cells (Agilent Technologies) and plated on LB with 100 µg/mL ampicillin and 10 µg/mL gentamycin. After overnight incubation at 30 °C, individual colonies were transferred to 5 mL LB/ampicillin and grown overnight at 30 °C. These cultures were used to inoculate 50 mL LB/ampicillin in 250 mL flasks (1:50 dilution) and grown at 30 °C to OD600 = 0.6–0.8. Cultures were cooled to 10 °C then protein expression induced by the addition of IPTG to 0.1 mM. Cultures were grown at 10 °C for 24 hours with shaking at 300 rpm. Cells were harvested (4000 rpm, 4 °C, 20 mins), resuspended in 1 mL Buffer A1 (50 mM HEPES pH 8.0, 50 mM glycine, 0.5 M NaCl, 30 mM imidazole, 5% v/v glycerol, EDTA free protease inhibitor tablet – 1 tablet/250 mL A1 buffer) and disrupted using a tissue homogenizer (Avestin) with a homogenizing pressure of 15,000 psi. By optimizing and adjusting the above basic methods, we also tried other different tags like SUMO, GB1–His, MBP–His in pOPIN–GG vector^14^ and different expression strains such as BL21 (DE3), Rosetta, SHuffle, Solu–BL21.

For large scale purification, (5 liters for *Pn*BAHD15), cell debris was removed by high-speed centrifugation at 45,000 × g for 45 mins. His-tagged protein was purified on a 5 mL Ni–NTA column. Purified protein was desalted and concentrated using a Vivaspin 30K centrifugal concentrator (Sartorius) according to manufacturer’s instructions. Protein was quantified by Nanodrop Microvolume Spectrophotometers (Thermo Fisher Scientific) and purity assessed by SDS–PAGE and InstantBlue Coomassie stain (Abcam).

### Assays of β-N-oxalyl-L-α,β-diaminopropionic acid synthase (β-ODAP synthase) activity using synthesised oxalyl-CoA

Assays were carried out in a total volume of 100 µL and were incubated at 30 °C for 1 hour. Purified BAHD15 enzyme was used at a concentration of 5–10 ng/µL and L-DAP at 2mM. Oxalyl-CoA concentrations varied from 0.1 mM–0.5 mM.

The velocity of the reaction was defined as the amount of ODAP produced in 1 hour. ODAP was measured, by derivatizing 20 µL samples of reaction mix using AccQ-Tag reagent (Waters, Milford, MA, USA) following the manufacturer’s instructions. Derivatized samples were diluted 1:100 (ODAP) in 0.1% (w/v) formic acid before LC–MS analysis. β-L-ODAP (Lathyrus Technologies, Hyderabad, India) standards were prepared and quantified using a Xevo triple quadrupole TQ–S instrument (Waters, Milford, MA, USA) as previously described. The amount of ODAP was defined as the area under the ODAP mass transition peak of 347>171.1. Km was determined from a Lineweaver–Burk Reciprocal plot.

Assays of purified PnBAHD15 showed that the enzyme could use oxalyl-CoA and L-DAP as substrates to synthesize β-ODAP. Kinetic analysis showed that the Km value of this reaction was 250 μM for oxalyl-CoA.

## Acknowledgements

We gratefully acknowledge support from the BBSRC ISP Grant “Harnessing Biosynthesis for Sustainable Food and Health (HBio)” (BB/X01097X/1) to the John Innes Centre.

## Supplementary Information

**Supplementary Figure 1.**
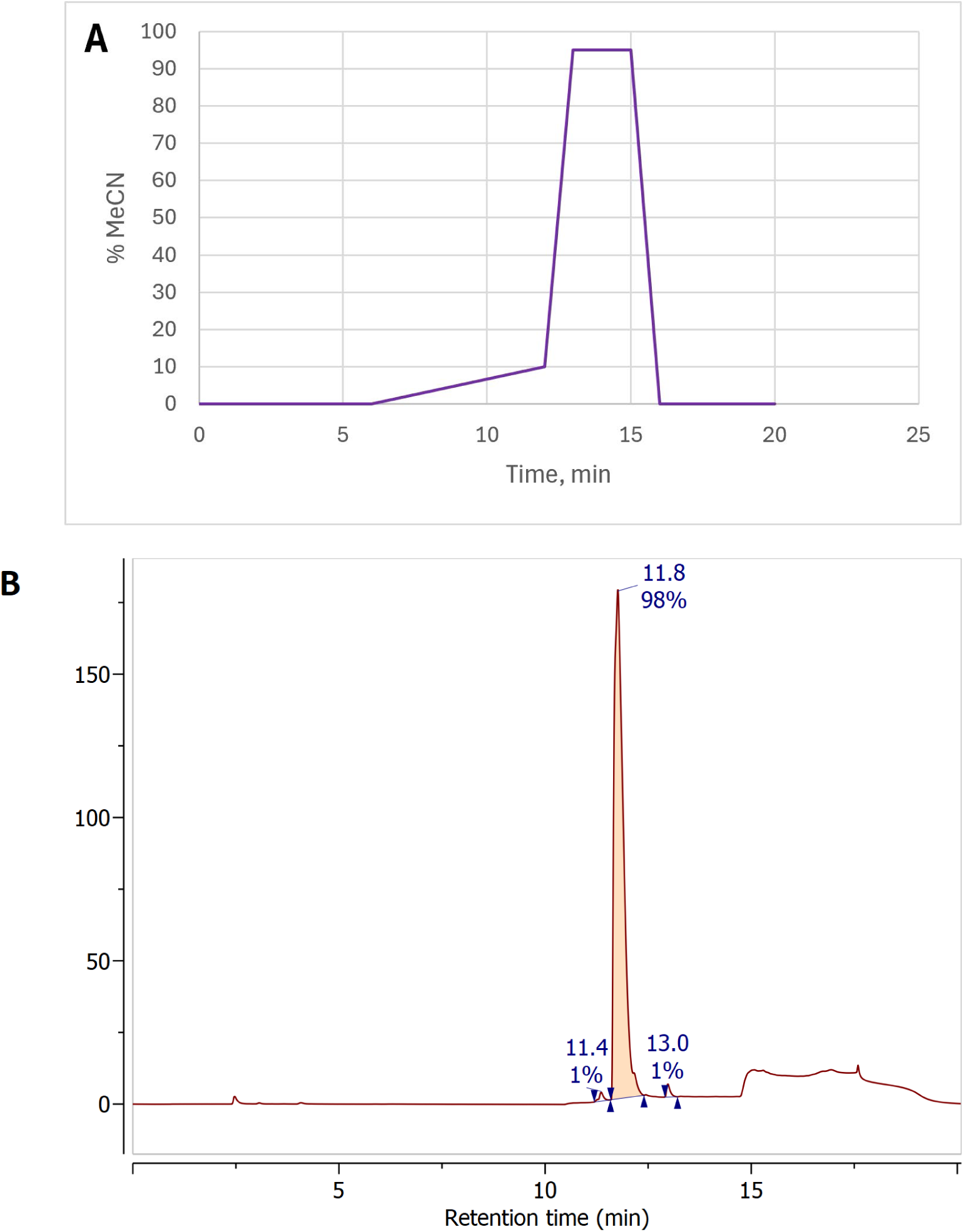
HPLC purification of oxalyl-CoA. (**A**) Elution method used for purification of oxalyl-CoA on a reverse-phase HPLC column (Phenomenex Kinetex, 5um XB-C18, 100 A, 250 x 21.2 mm) using gradient of MeCN in 10 mM ammonium formate. (**B**) Elution profile of products of oxalyl-CoA synthesis. Note that peak of unmodified CoA was not detected. In the applied conditions CoA elutes 1.5 min after oxalyl-CoA (data not shown).

**Supplementary Figure 2.**
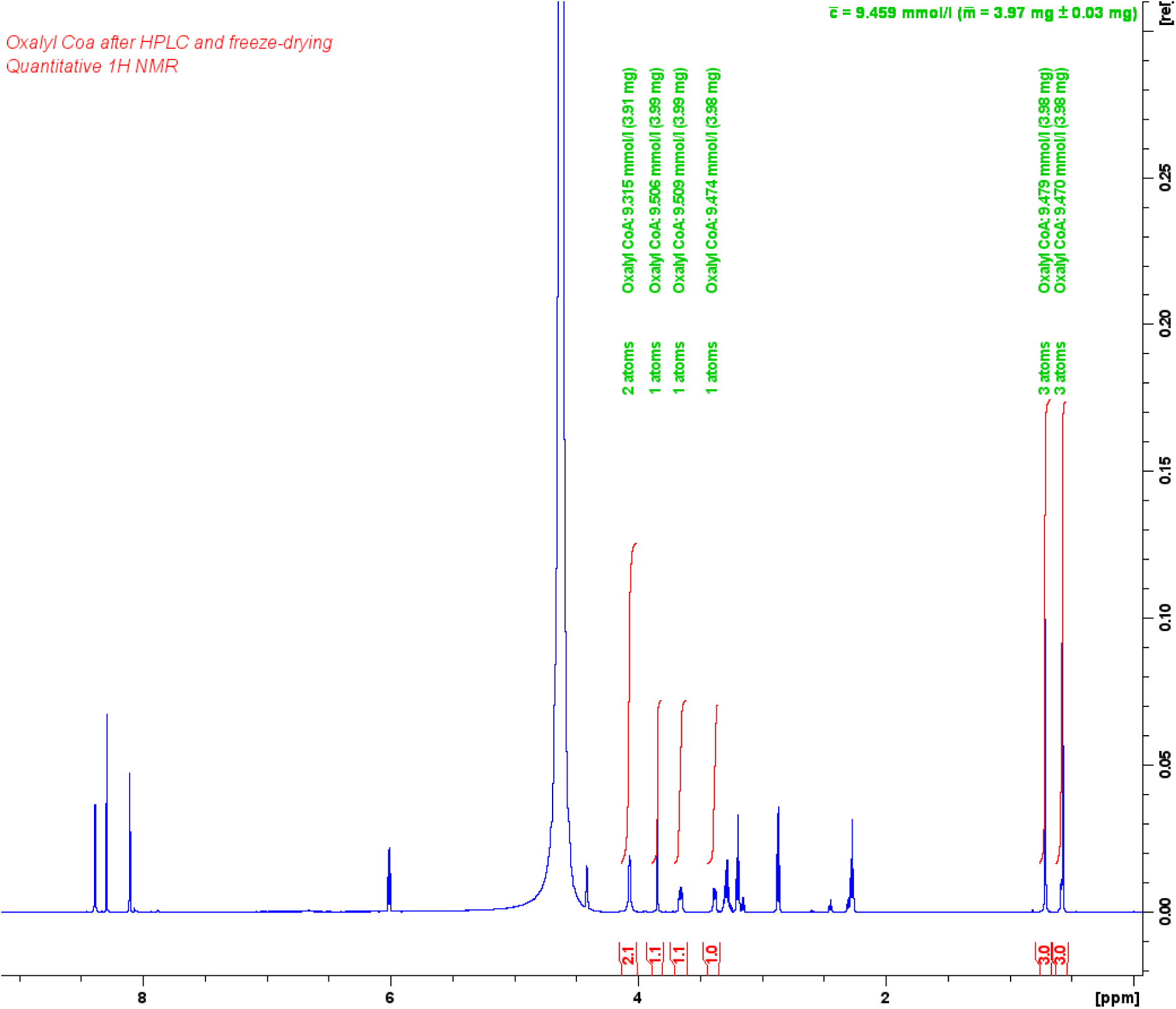
Quantification of purified oxalyl-CoA by qHNMR. The amount of purified oxalyl-CoA was determined by quantitative 1H NMR spectroscopy in D2O using the TopSpin ERETIC protocol and calculated as 4.0 mg. ERETIC calibration was performed using 10 mM dimethyl sulfone in D2O as an external standard.

**Supplementary Figure 3.**
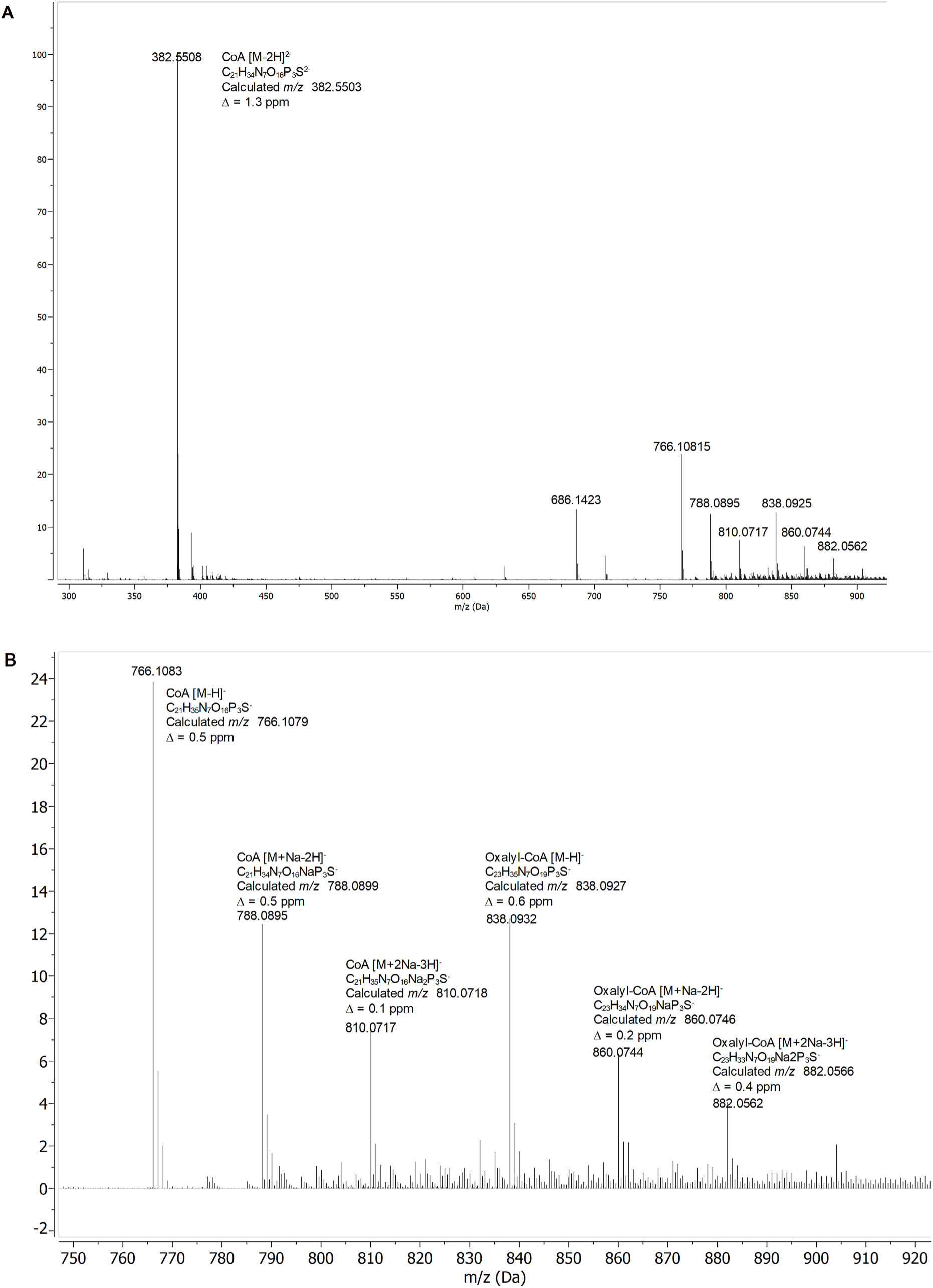
HR-ESI-MS spectra of purified of oxalyl-CoA (**3**). Spectra were acquired in negative mode. (**A**) Full mass spectrum *m/z* range 300-950 and (**B**) annotated region of the spectrum (*m/z* 750-920). The ESI-MS spectrum of oxalyl-CoA was characterised by the presence of ions corresponding to [M−H]^−^, [M+Na−2H]^−^ and [M+2Na−3H]^−^. Under the MS conditions, cleavage of the thioester bond resulted in the formation of CoA-derived ions with analogous charge states. The most intense signal corresponded to the doubly charged [M−2H]^2−^ ion of CoA.

**Supplementary Figure 4.**
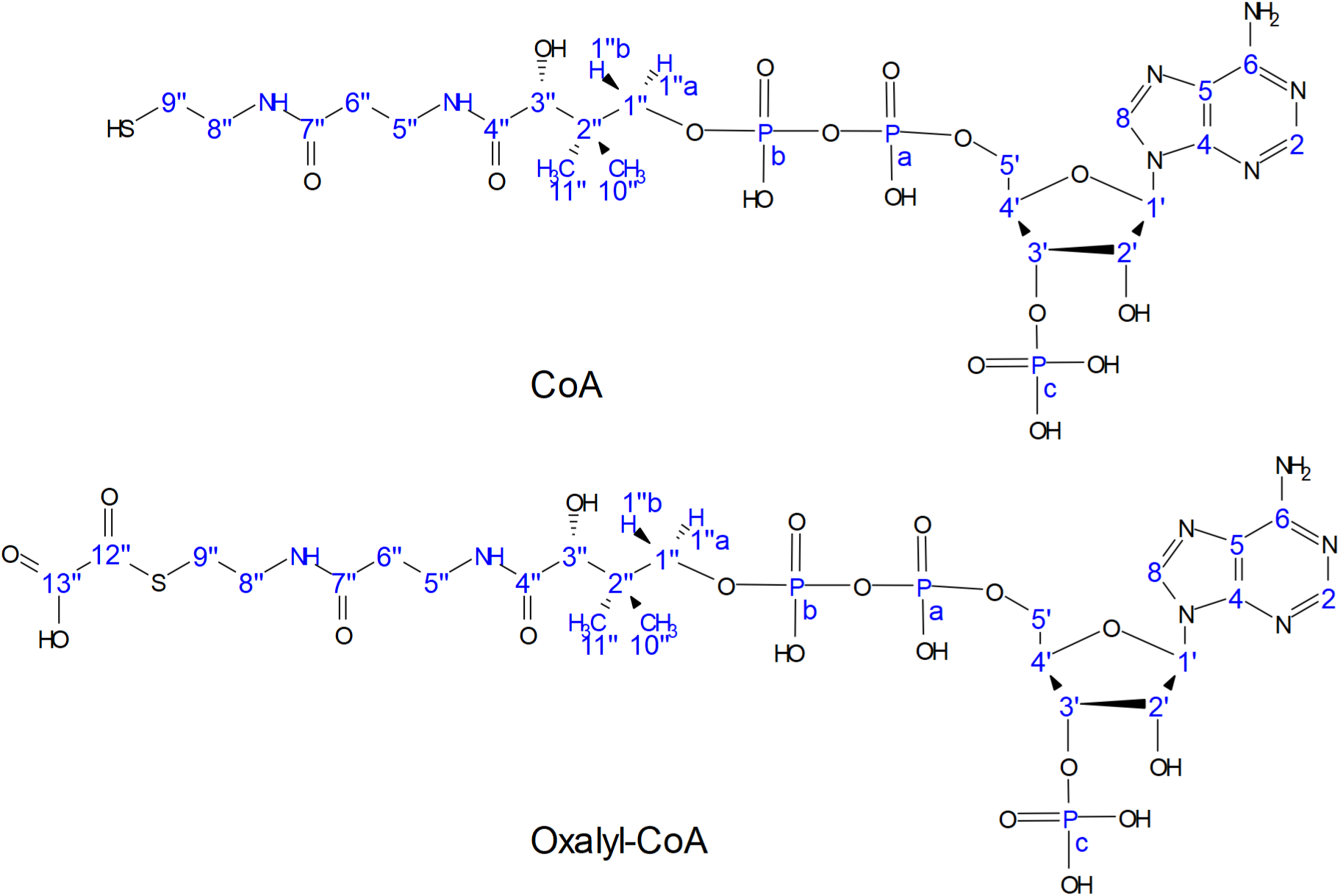
Structures of CoA and oxalyl-CoA. Atom numbering used for NMR signal assignment is indicated.

**Supplementary Figure 5.**
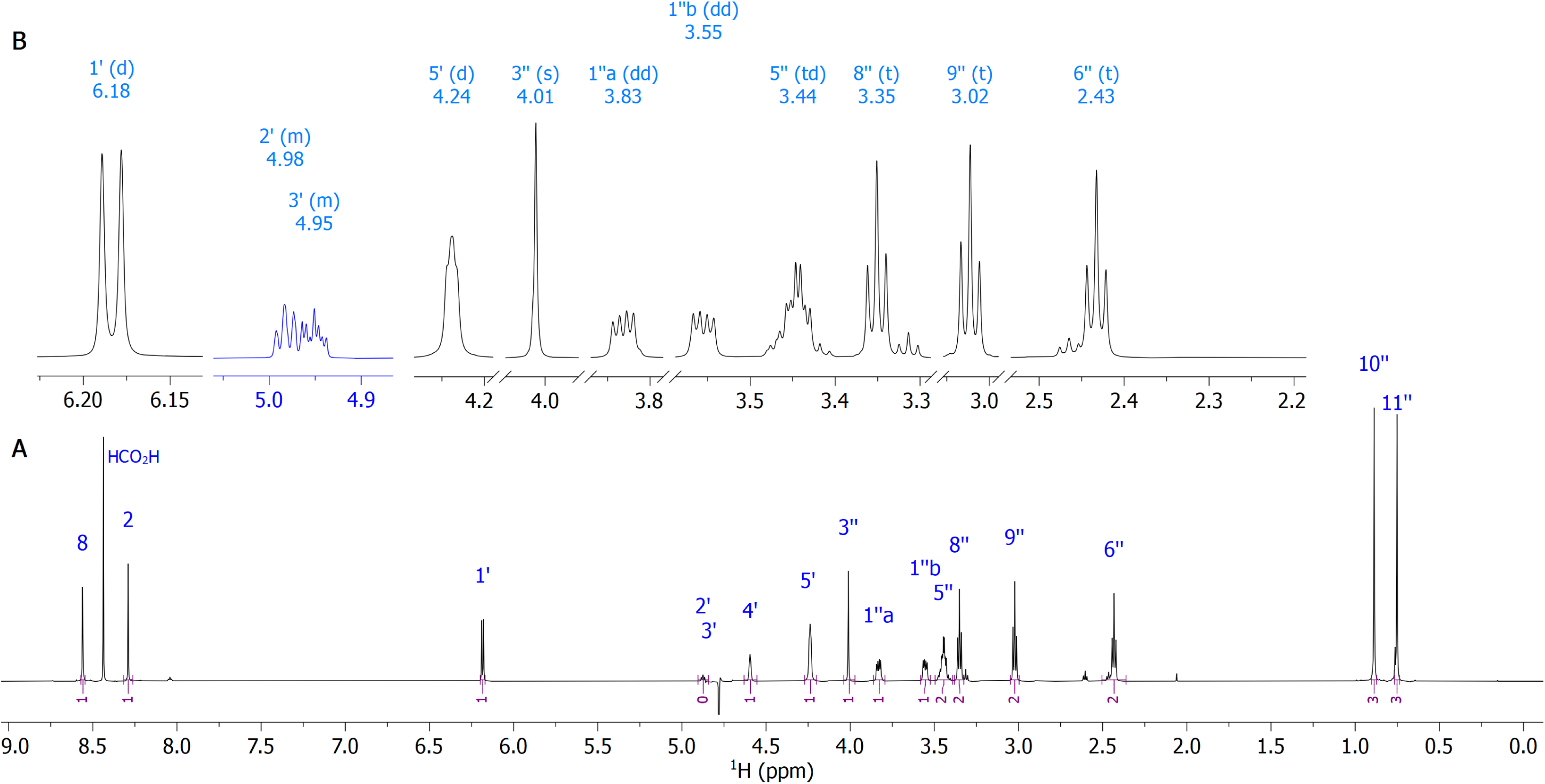
^1^H NMR spectra of oxalyl-CoA. The spectra were recorded with water suppression by presaturation. (**A**) Full spectrum overview; singlet at δ 8.44 ppm corresponds to residual formate from the HPLC buffer. (**B**) Expanded region showing multiplets. The fragment between δ 4.9–5.5 ppm was taken from a spectrum recorded at 308 K to reveal signals of H-2’ and H-3’, which were obscured by the residual water signal at 298 K.

**Supplementary Figure 6.**
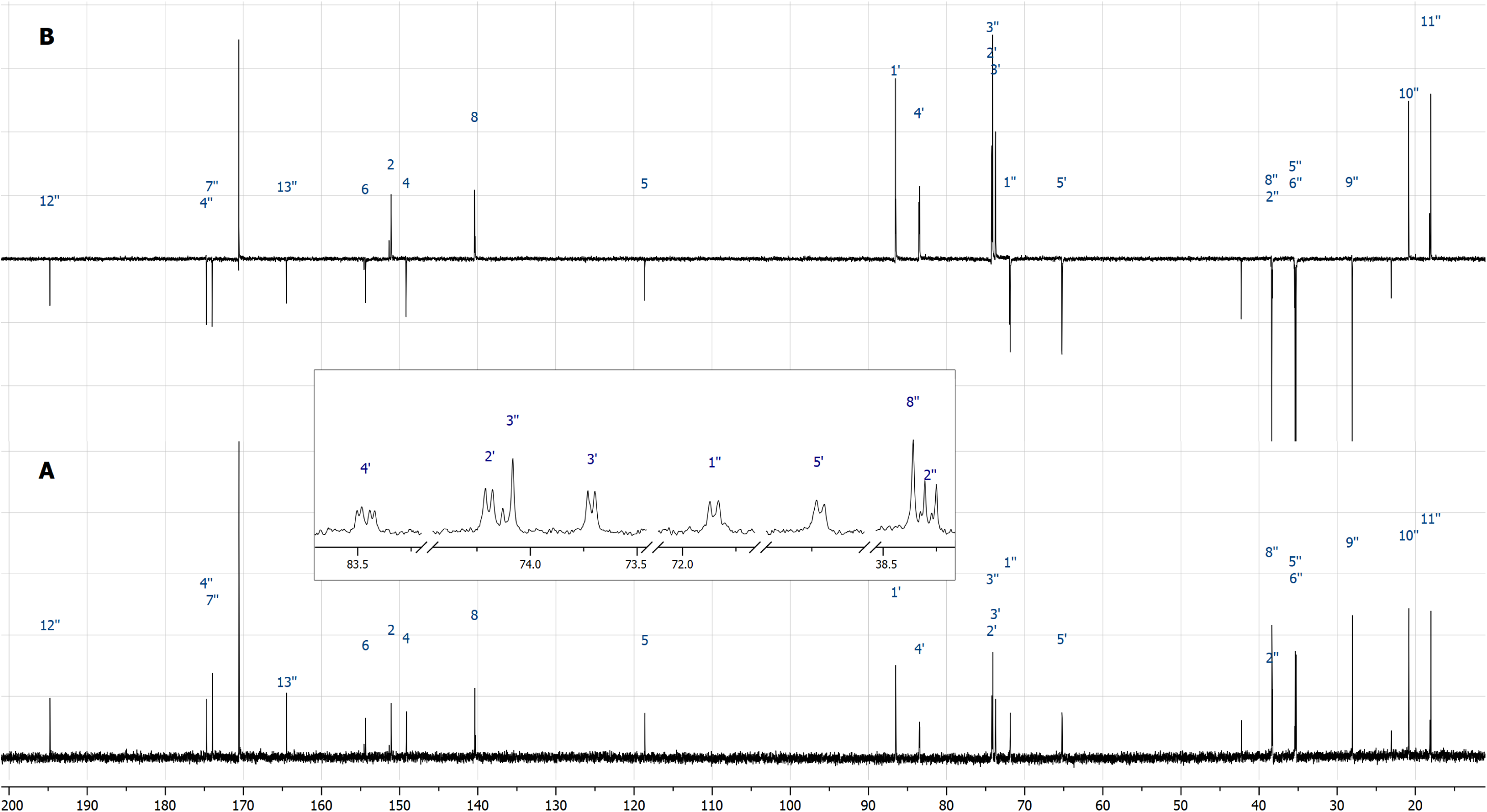
Overlay of ^13^C NMR and DEPTQ spectra of oxalyl-CoA. (**A**) ^13^C NMR spectrum and (**B**) DEPTQ spectrum. The insert shows an expansion containing ^13^C resonances split as a result of ^13^C–^31^P couplings.

**Supplementary Figure 7.**
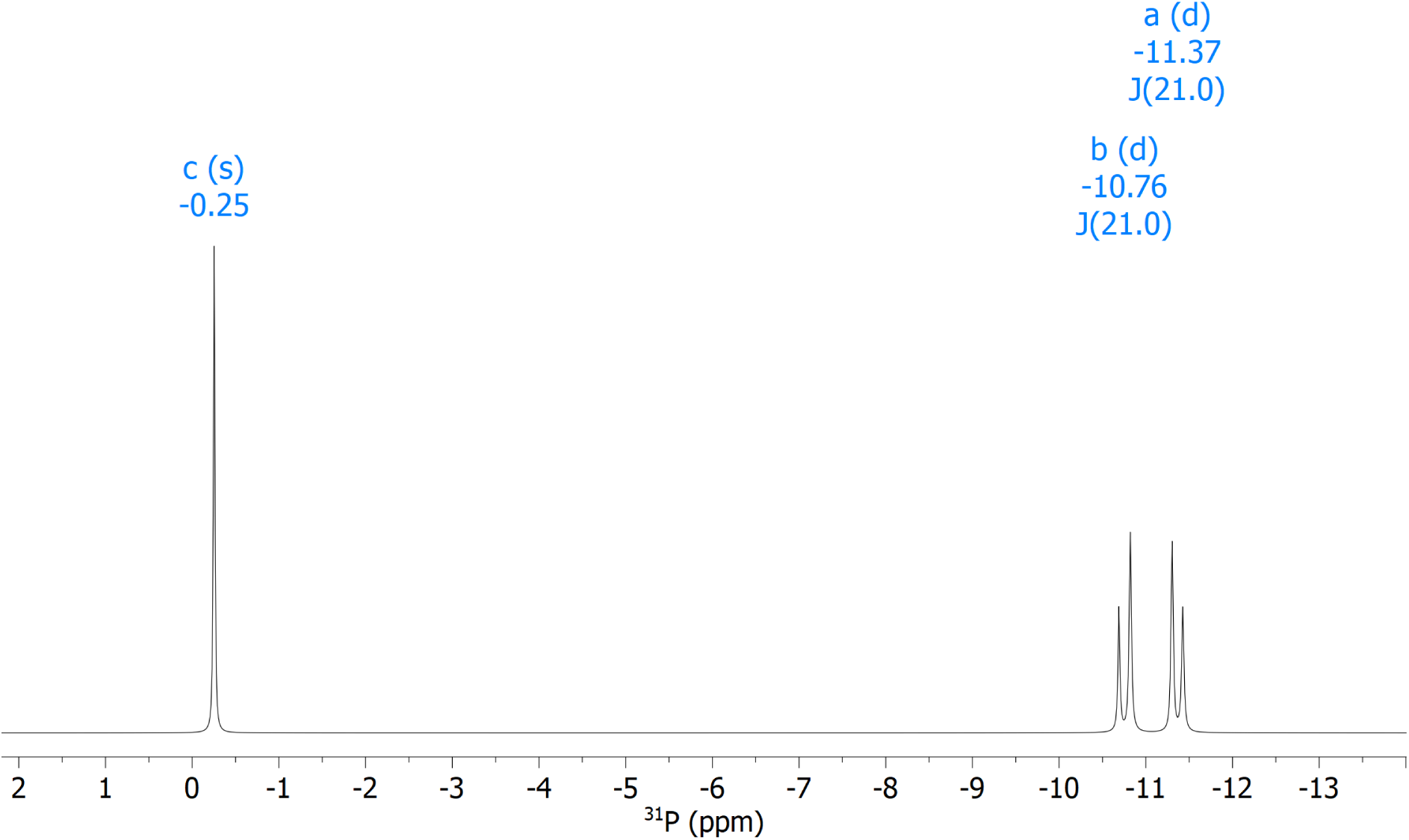
^31^P NMR (162 MHz) spectrum of oxalyl-CoA.

**Supplementary Figure 8.**
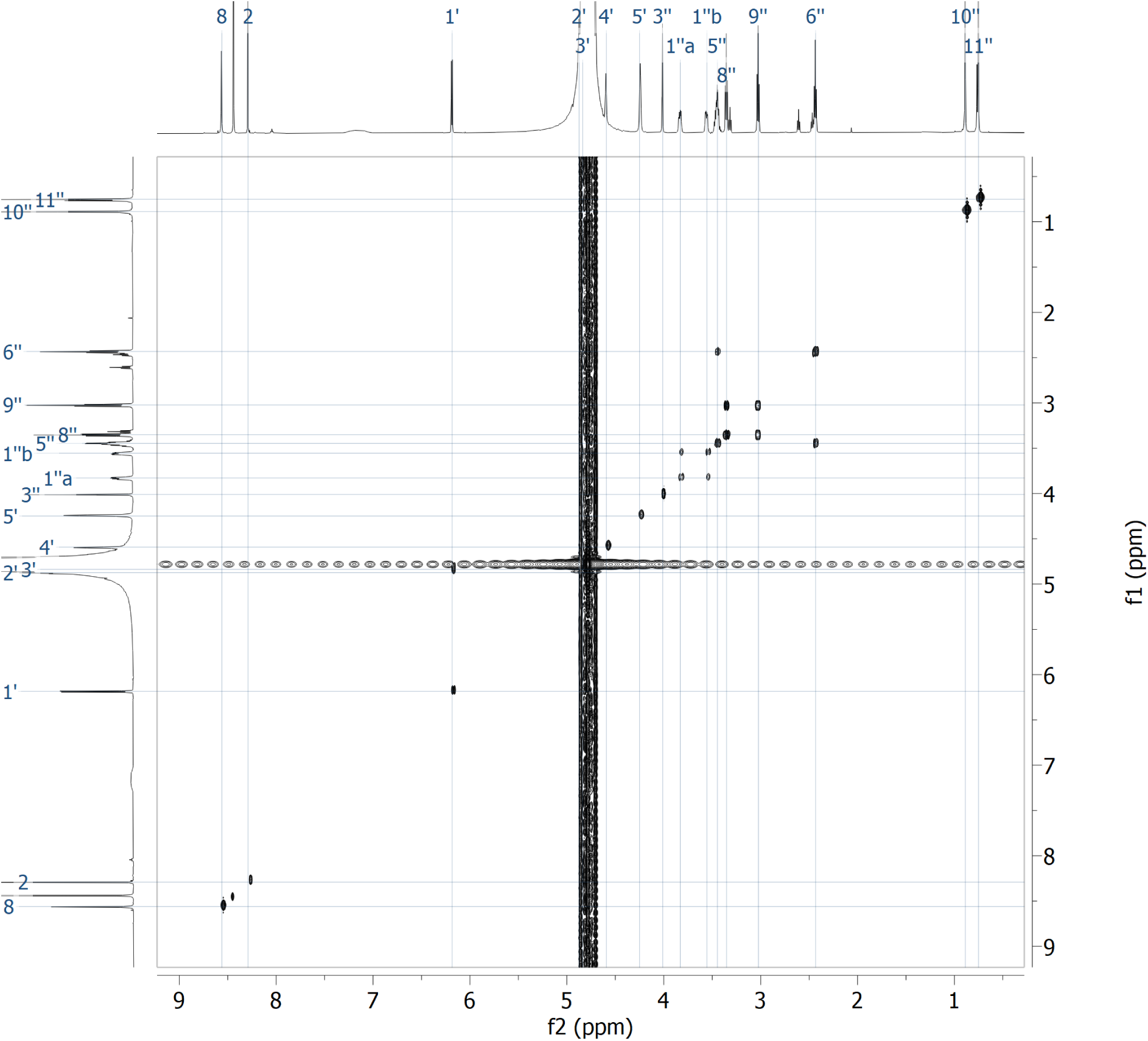
^1^H,^1^H-COSY NMR spectrum of oxalyl-CoA.

**Supplementary Figure 9.**
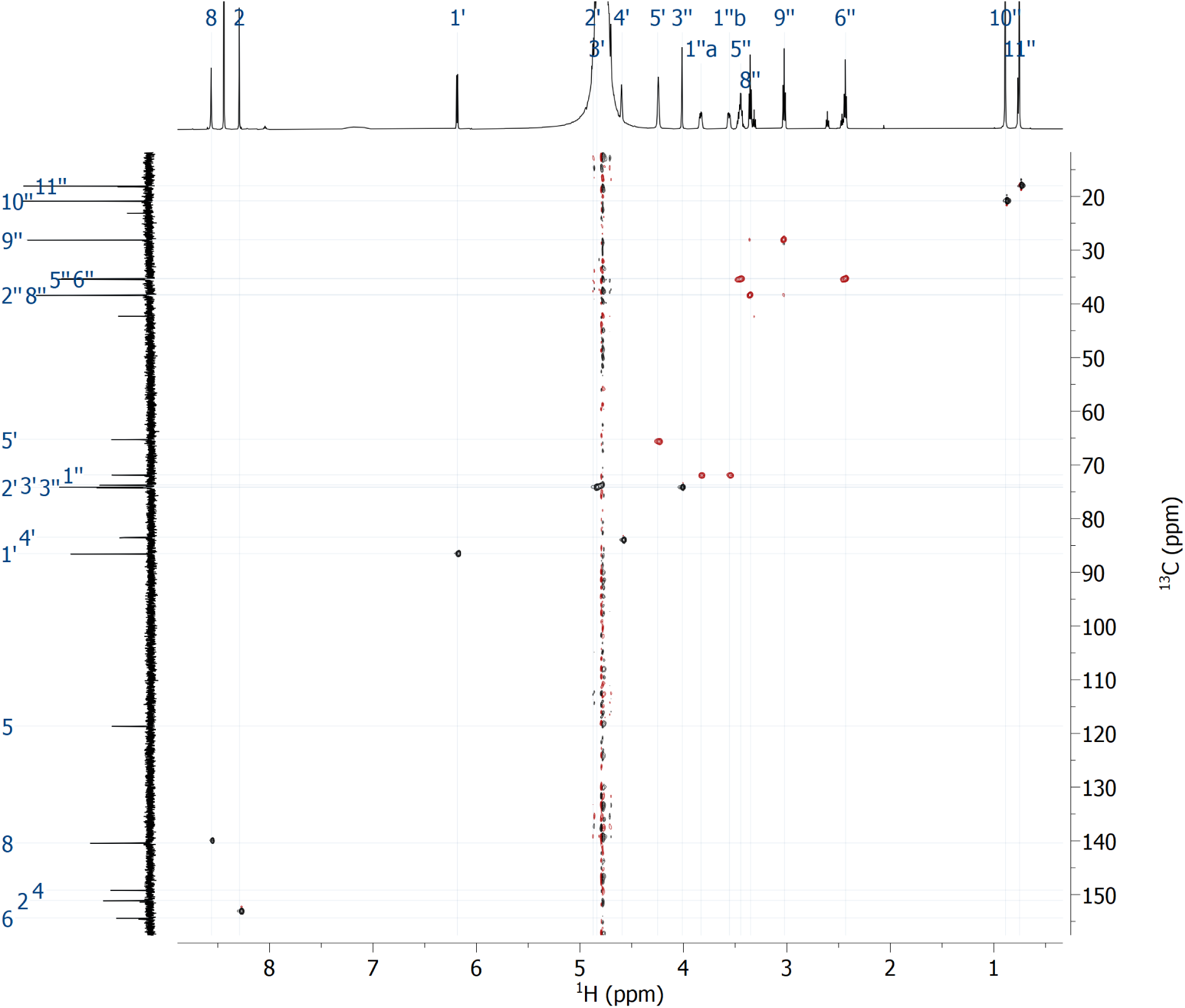
^1^H,^13^C-HSQC NMR spectrum of oxalyl-CoA.

**Supplementary Figure 10.**
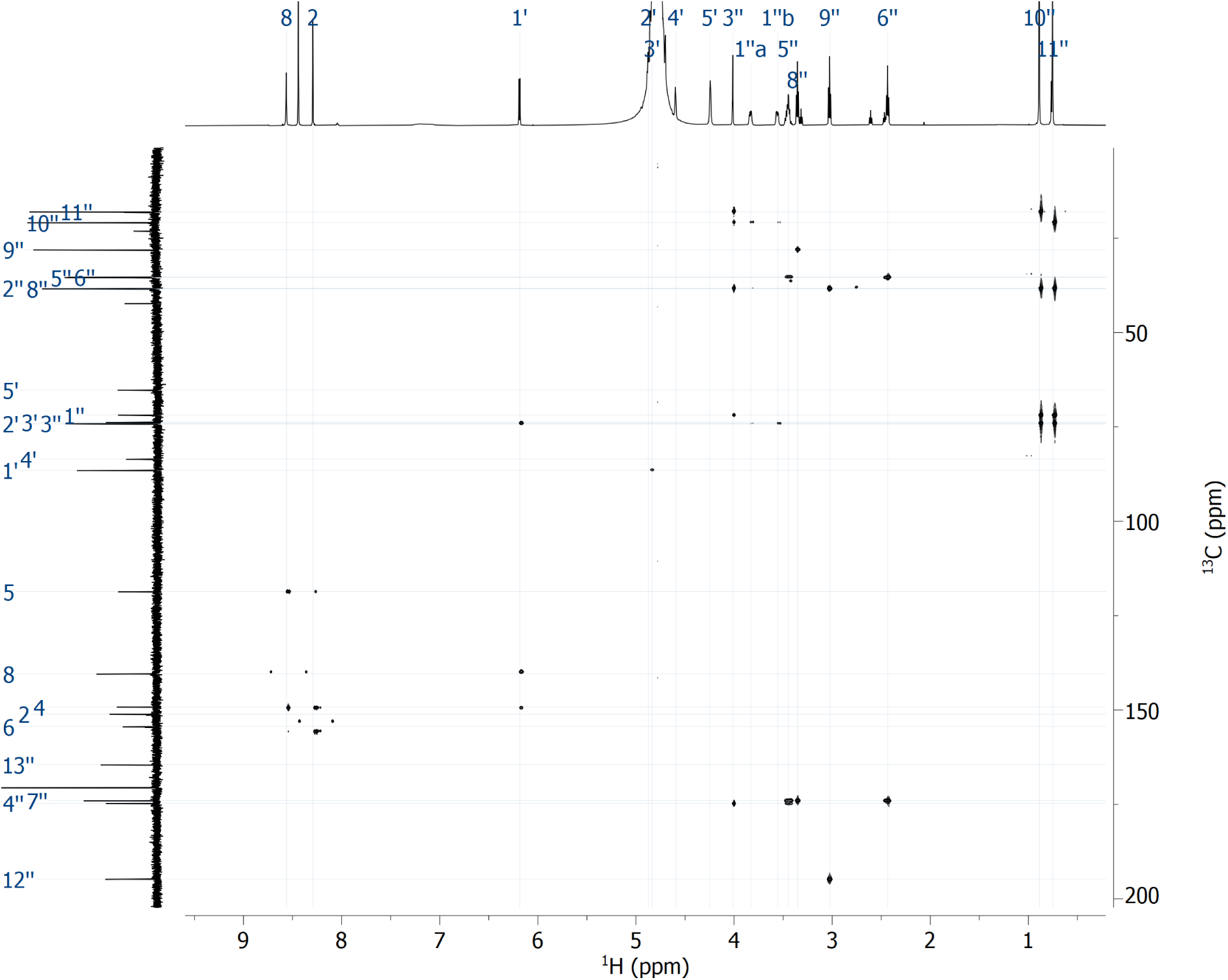
^1^H,^13^C-HMBC NMR spectrum of oxalyl-CoA.

**Supplementary Figure 11.**
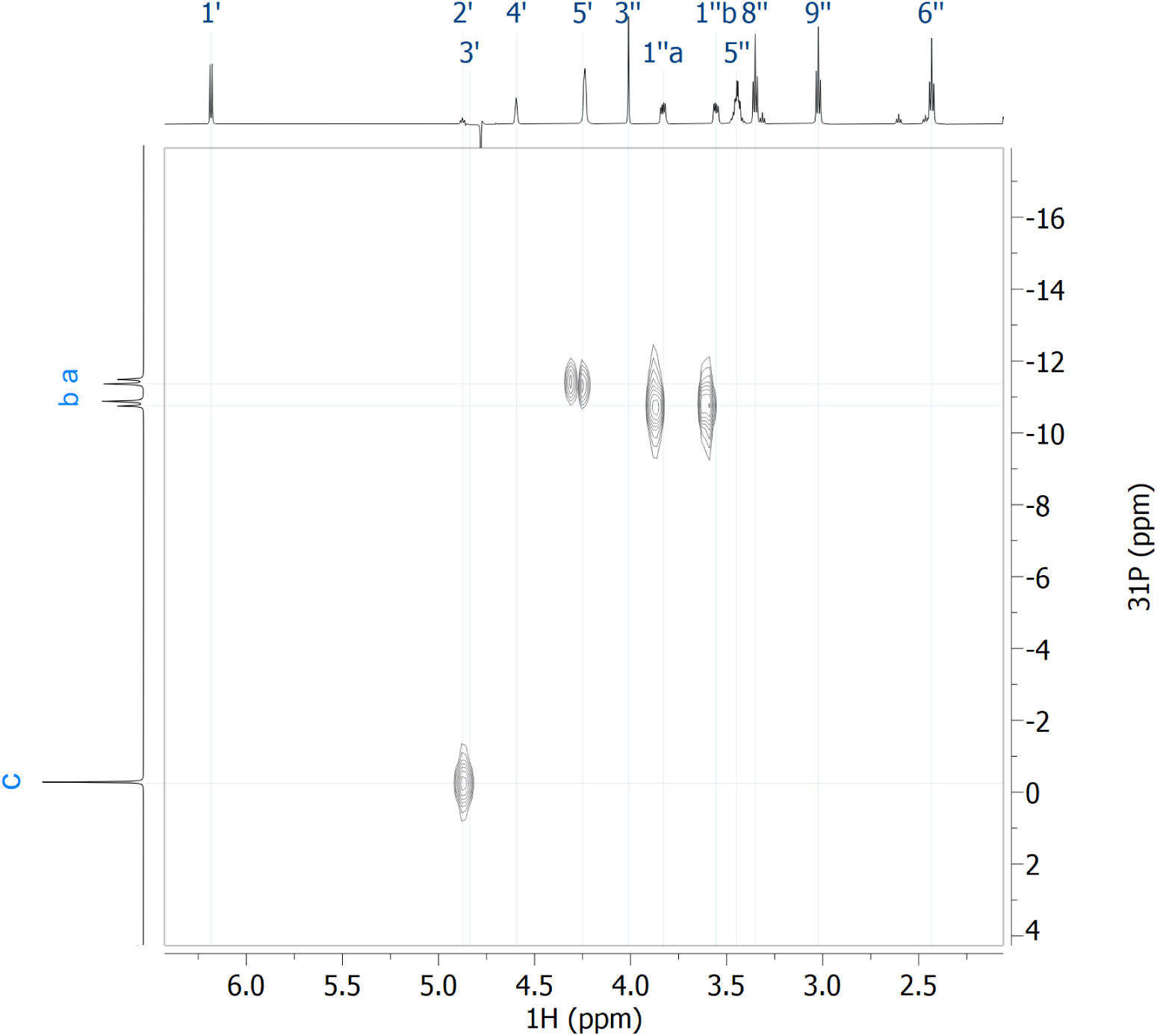
^1^H,^31^P-HMBC NMR (400 MHz) spectrum of oxalyl-CoA.

**Supplementary Figure 12.**
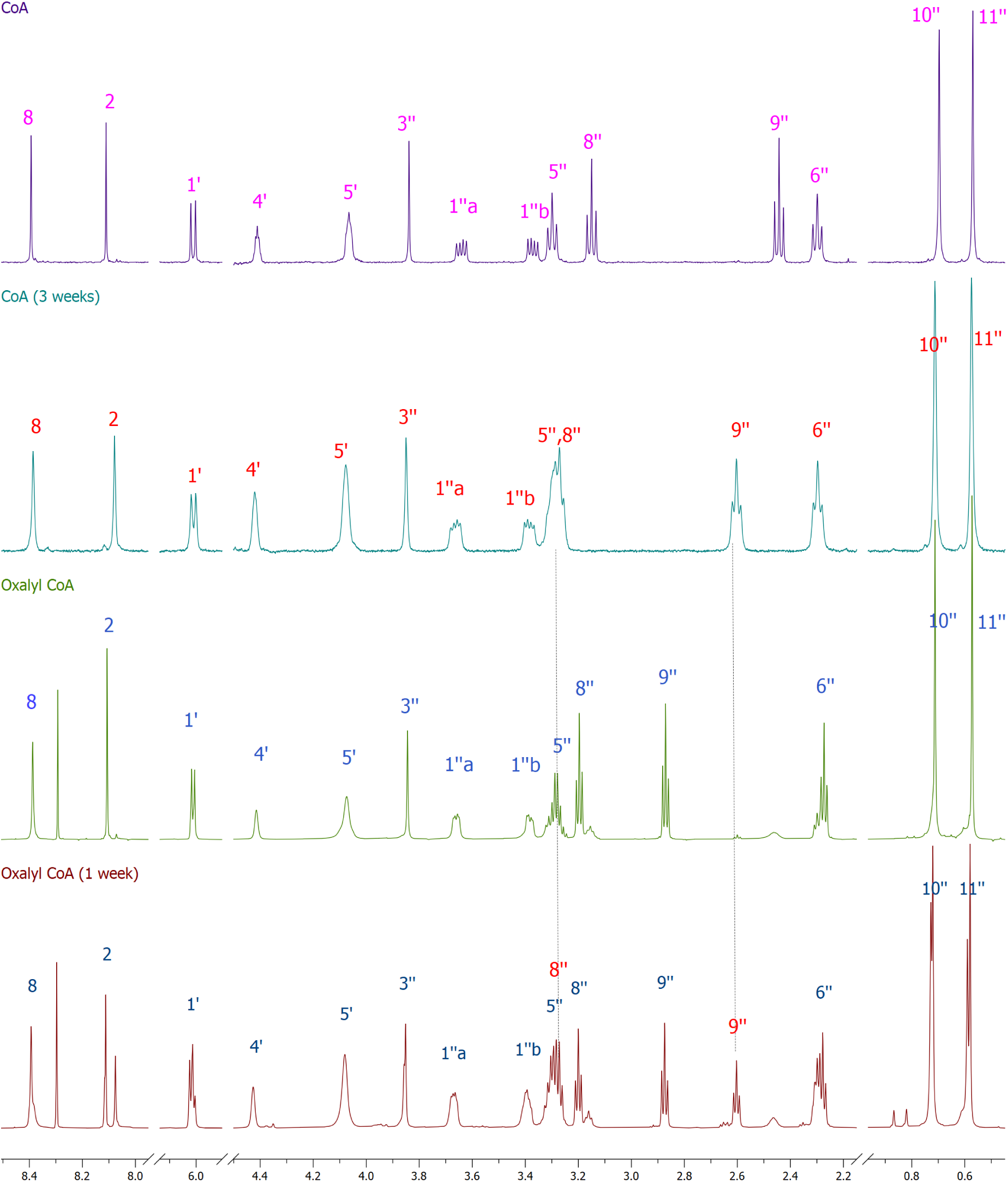
Degradation of CoA and oxalyl-CoA in a D_2_O during storage. Stack of assigned ^1^H NMR spectra (D_2_O, 298 K) of freshly prepared CoA, the same sample CoA sample after storage at 4 °C for 3 weeks, freshly prepared oxalyl-CoA and the sample of oxalyl-CoA stored at 4 °C for 1 week. Atom numbering used for assignment of signals is shown in Supplementary Fig. 4.

